# MembraneFold: Visualising transmembrane protein structure and topology

**DOI:** 10.1101/2022.12.06.518085

**Authors:** Santiago Gutierrez, Wojciech G. Tyczynski, Wouter Boomsma, Felix Teufel, Ole Winther

**Affiliations:** DTU Compute, Technical University of Denmark, Lyngby, Denmark; Department of Computer Science, University of Copenhagen, Copenhagen, Denmark; Department of Biology, University of Copenhagen, Copenhagen, Denmark; Digital Science and Innovation, Novo Nordisk A/S, Måløv, Denmark; Genomic Medicine, Rigshospitalet, Copenhagen, Denmark

**Keywords:** Transmembrane proteins, Protein structure, Transmembrane protein topology

## Abstract

**Background:** AlphaFold’s accuracy, which is often comparable to that of experimentally determined structures, has revolutionized protein structure research. Being a statistical method, AlphaFold implicitly infers the cellular environment, e.g. the cell membrane, from the protein sequence. Membrane protein topology prediction methods predict the cellular environment for each protein residue but not the structure. Current structure and topology tools thus provide complementary information.

**Results:** We introduce the web server MembraneFold. MembraneFold combines protein structure (from an uploaded PDB file/AlphaFold DB/OmegaFold) and topology (DeepTMHMM) prediction in one server. The output is shown both as a structure with topology superimposed and as a sequence annotation. MembraneFold uses structures predicted by OmegaFold if neither a PDB file is uploaded nor the structure is available in AlphaFold DB.

**Conclusion:** MembraneFold is a user-friendly web server that provides practitioners with fast and accurate information about membrane proteins. It is available at https://ku.biolib.com/MembraneFold/.

## Background

AlphaFold DB (`alphafold.ebi.ac.uk`) provides predicted structures for all UniProt sequences, making it an invaluable resource for protein structure research. For sequences not in UniProt, it is possible to obtain structure predictions by running AlphaFold [1] and emerging alternatives such as OmegaFold [2].

The current approach to protein structure prediction is based upon using the protein sequence information only. It can predict structures of membrane as well as globular proteins including those that rely on other molecules to stabilize their structure (such as Zinc finger proteins) [3]. The cellular environment is thus implicitly inferred from the sequence information. In the case of protein multimers and transmembrane proteins we are interested in predicting how the protein interacts with other proteins and the membrane, respectively. AlphaFold has been extended to multimers [4] and placement of the protein in the membrane can be predicted with so-called transmembrane topology methods, e.g. [5, 6, 7, 8, 9].

In this paper we present a web server MembraneFold (ku.biolib.com/MembraneFold) that combines structure and topology prediction in one shared view. The workflow of MembraneFold is shown in Figure 1. The user may either provide sequence or structure information (in fasta or PDB format). If the sequence information is provided, then the structure is retrieved from AlphaFold DB, and if it is not available there, OmegaFold is used for structure prediction [2]. OmegaFold is a faster alternative to AlphaFold with comparable performance [2].

**Figure 1:**
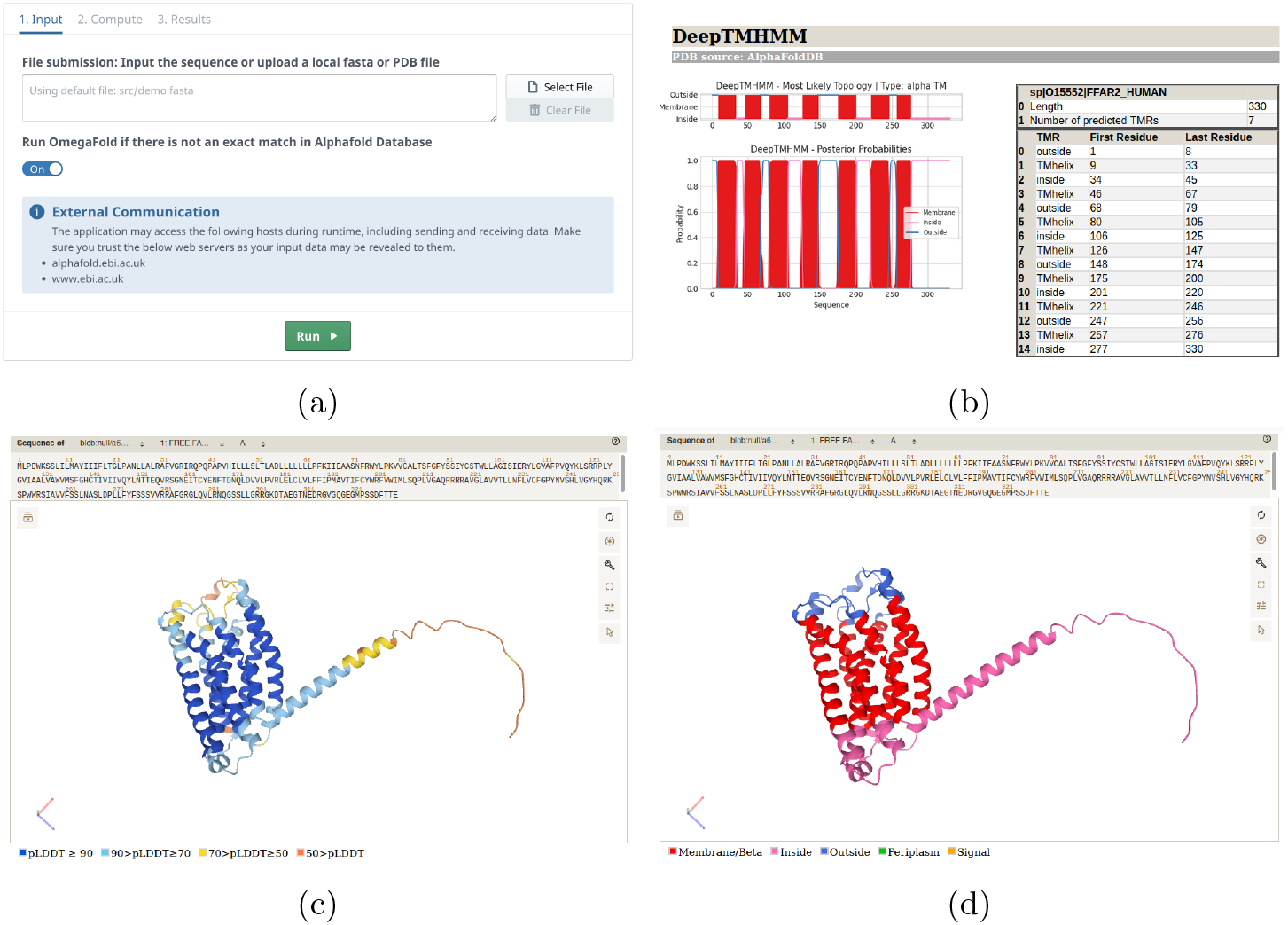
MembraneFold overview. (a) The input is a fasta sequence/file or a PDB structure. For fasta input, MembraneFold performs a look-up in https://alphafold.ebi.ac.uk/ and if not present predicts the structure with OmegaFold [2]. MembraneFold outputs the structure with (b) the DeepTMHMM topology superimposed on the structure, (c) pLDDT confidence annotation if the structure is predicted and B-factor if the structure is experimental and (d) DeepTMHMM topology sequence format.

### MembraneFold

MembraneFold combines two types of protein sequence prediction tools: protein structure (AlphaFold [1] and OmegaFold [2]) and transmembrane protein topology (DeepTMHMM [5]). The Mol* toolkit [10] is used for visualisation.

DeepTMHMM is a protein language model based neural network that predicts the transmembrane (TM) topology of proteins directly from their sequence. For each protein, it predicts a protein type label: Alpha (helical) TM, alpha TM with Signal peptide (SP), beta barrel TM, globular and globular with SP. For each residue, it also predicts a label: Signal peptide (S), inside cell/cytosol (I), alpha membrane (M), beta membrane (B), periplasm (P), and extracellular/lumen of ER/Golgi/lysosomes (O) [5].

AlphaFold [1] is a machine learning model that predicts the 3D structure of a protein based on its amino acid sequence. AlphaFold was, by a wide margin, the top-ranked protein structure prediction method in CASP14 (Critical Assessment of protein Structure Prediction), with high-accuracy predictions. While AlphaFold occasionally struggles with disordered regions, the CASP findings and the widespread use of AlphaFold imply that AlphaFold is becoming an invaluable tool for protein research.

The purpose of MembraneFold is to present membrane positioning and structure predictions of a given protein chain simultaneously. The tool allows to obtain a fast overview of this information and quickly toggle between the topology annotation and the structure confidence scores.

As illustrated in Figure 2, MembraneFold has three main modules: topology prediction, structure prediction or retrieval and visualization. The tool can handle one protein at a time and can either be used through the web-interface or as as command line tool. It starts with parsing a sequence from the user input. The input can be introduced in different formats: PDB file, fasta file or if using the web-interface a string input.

**Figure 2:**
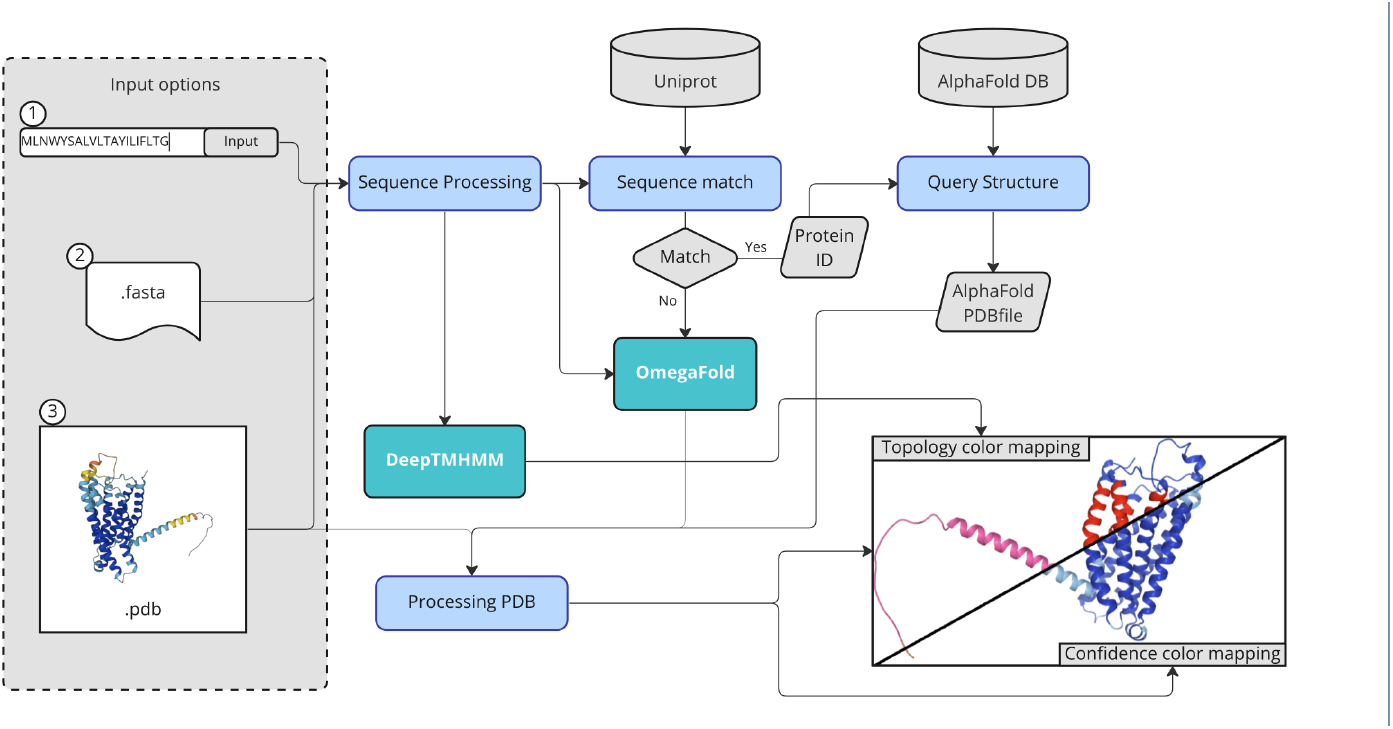
Overview of MembraneFold.

The first module initializes and runs DeepTMHMM which produces the topology prediction of the given sequence. MembraneFold is meant to work only with membrane proteins, therefore, if the prediction of DeepTMHMM does not contain membrane sections, the computation stops, a notification is displayed to the user and the option to download the DeepTMHMM prediction is provided.

After obtaining the membrane positioning prediction, the structure module is initialized. This module is in charge of obtaining the 3D structure of the protein. Firstly, if a PDB file is provided then this is used for all the downstream analysis. If a sequence is provided, we look for an exact sequence match in the Uniprot database. If the query is successful, MembraneFold extracts the correspoding structure from the AlphaFold database. Otherwise, the protein structure is predicted using OmegaFold.

Once both predictions are received, the processing and visualization part creates the front-end layer of the tool. The user interface output page consists of two tabs: DeepTMHMM and pLDDT Confidence, or B-factors if the structure is experimental. In both tabs, the 3D structure visualization of the amino acid chain, together with the colour mask over it and information about the source of the PDB file are displayed. In the pLDDT Confidence tab, the colour of the 3D model indicates the predicted local distance difference test score (pLDDT score) of the amino acids along the chain. The pLDDT ranges between 0 and 100 and indicates how well the prediction would match an experimental structure. This confidence is based on the local distance difference test C*α* (lDDT-C*α*). In the case of the DeepTMHMM tab, the colour on the structure visualization maps the topology of each residue to the 3D structure. In addition, Further below in the topology tab, it is presented statistical graphs obtained from DeepTMHMM showing the probabilities of the prediction and a table with description of the topology per section. MembraneFold output files (plots, PDB and fasta files) are downloadable in one zip file.

### Related work

Historically, the prediction of transmembrane topology from sequence was motivated by the unavailability of structural information, as transmembrane proteins are notoriously difficult to crystallize [11]. However, once the structure of a transmembrane protein becomes known, the exact membrane positioning still needs to be inferred computationally, as its experimental determination is very costly [12]. To this end, physics-based approaches such as PPM [12] (opm.phar.umich.edu/ppm_ server) and TMDET [13] were developed. The underlying principle is the insertion of the protein structure into a lipid bilayer so that the energy of the system is optimized. This is either done using solvation models in PPM or a simplified objective function in TMDET. While these approaches are capable of accurately determining the membrane-buried residues, they require the user to specify the topology of the N-terminus, as they cannot differentiate between outside and inside of the membrane. Furthermore, they can suffer from AlphaFold’s spurious folding of disordered regions and signal peptides that can violate membrane constraints [14].

The availability of predicted structures has spawned the expansion of expert-curated resources such as Membranome [15] (`membranome.org`), which offer comprehensive information about the membrane topology of proteins and take advantage of the mentioned structural insertion algorithms to provide visualizations of proteins in their membrane context. However, given the nature of expert curation, they offer no easy way for the user to directly investigate a transmembrane protein from the sequence only. Another resource is TmAlphaFold [14] that has more than 200 thousand AlphaFold structures predicted to be alpha helical transmembrane proteins by CCTOP [7] with membrane placement predicted by TMDET [13].

The supplementary material showcases data analysis with MemebraneFold for a number of proteins with links to the corresponding models in Membranome and TmAlphaFold when available.

## Conclusion

MembraneFold is a simple and easy-to-use tool that visualises transmembrane protein structures with their membrane context. Before the availability of MembraneFold there was no single place to go when wanting to superimpose predicted transmembrane protein topology on the structure. The tool is highly modular so when new structure and topology prediction methods becoms available, these can easily be integrated into MembraneFold.

## Acknowledgements

We want to thank Frederikke I. Marin, Marloes Arts and Jonah Tabbal for many useful discussions and the BioLib team for software help.

## Funding

W.B., F.T. and O.W. are supported by the Novo Nordisk Foundation (NNF20OC0062606).

## Availability of data and materials

Project name: MembraneFold

Project home page: https://ku.biolib.com/MembraneFold/

Operating system(s): Platform independent

Programming language: Python

Other requirements: No other installations required. Uses DeepTMHMM, AlphaFold DB and Mol*.

Licenses: Provided as is. Any restrictions to use by non-academics: The server is made freely available here for both academic and commercial purposes. For commercial users wishing to run the software on their own servers, DeepTMHMM requires a commercial license.

## Ethics approval and consent to participate

Not applicable

## Competing interests

The authors declare that they have no competing interests.

## Consent for publication

Not applicable

## Authors’ contributions

The authors together designed MembraneFold. S.G. and W.G.T. - equal contributions - implemented MembraneFold. S.G., W.G.T., F.T. and O.W. wrote the paper. All authors have read and approved the final version of the paper.

## Notes

https://biolib.com/KU/MembraneFold/

### Competing Interest Statement

The authors have declared no competing interest.

