## supplementary material for "MembraneFold: Visualising transmembrane protein structure and topology"

### MembraneFold supplementary data analysis

We provide data analysis of a small set of proteins with MembraneFold and comparative tools: Membranome and TmAlphaFold.

For the investigated proteins listed below there were no hits in the Membranome database and only a few hits in TmAlphaFold.

#### Proteins investigated

The table below lists proteins investigated, their type, the source of structure and sequence and link to the protein in TmAlphaFold if available. All MembraneFold results including those presented in the example section below are available in this folder https://drive.google.com/drive/folders/1dyGL1otbCB2aGQ4-e_FEvJskhp4mYvjQ?usp=sharing.

| Protein | Type | Structure | Sequence | Reference | TmAlphaFold |
|---|---|---|---|---|---|
| Q9UBN4 | Alpha TM | https://alphafold.ebi.ac.uk/entry/Q9UBN4 | https://alphafold.ebi.ac.uk/entry/Q9UBN4 | https://www.ncbi.nlm.nih.gov/pmc/articles/PMC8761152/ | https://tmalphafold.ttk.hu/entry/Q9UBN4 |
| Q48658 | Alpha TM | https://alphafold.ebi.ac.uk/entry/Q48658 | https://www.uniprot.org/uniprotkb/Q48658/entry | https://www.ncbi.nlm.nih.gov/pmc/articles/PMC8761152/ | N/A |
| A0A7L3KCZ9 | Alpha TM | https://alphafold.ebi.ac.uk/entry/A0A7L3KCZ9 | https://alphafold.ebi.ac.uk/entry/A0A7L3KCZ9 | https://www.ncbi.nlm.nih.gov/pmc/articles/PMC8761152/ | N/A |
| P64606 | AlphaTM | https://www.rcsb.org/structure/7DUW | https://www.rcsb.org/structure/7DUW | https://www.ncbi.nlm.nih.gov/pmc/articles/PMC8761152/ | https://tmalphafold.ttk.hu/entry/P64606 |
| A0A4W2BVX4 | Alpha TM | https://alphafold.ebi.ac.uk/entry/A0A4W | https://alphafold.ebi.ac.uk/entry/A0A4W | Search AF DB for "transmembr | N/A |

| | | | | | |
|---|---|---|---|---|---|
| | | 2BVX4 | 2BVX4 | ane” | |
| A0A7J7Y2V0 | Alpha TM | https://alphafold.ebi.ac.uk/entry/A0A7J7Y2V0 | https://alphafold.ebi.ac.uk/entry/A0A7J7Y2V0 | Search AF DB for “transmembrane” | N/A |
| L8XZM1 | AlphaTM | https://alphafold.ebi.ac.uk/entry/L8XZM1 | https://alphafold.ebi.ac.uk/entry/L8XZM1 | https://pubmed.ncbi.nlm.nih.gov/23385571/ | N/A |
| P42055 | Beta barrel TM | https://alphafold.ebi.ac.uk/entry/P42055 | https://alphafold.ebi.ac.uk/entry/P42055 | https://pubs.acs.org/doi/10.1021/acs.jproteome.9b00740 | N/A |
| Q7SBE0 | Beta barrel TM (Uniprot) and Globular DeepTMHMM | https://alphafold.ebi.ac.uk/entry/Q7SBE0 | https://www.uniprot.org/uniprotkb/A0A7J7Y2V0/entry | AF DB structurally similar (FoldSeek) to P42055 | N/A |
| S3CXT0 | Beta barrel TM (Uniprot) and Globular DeepTMHMM | https://alphafold.ebi.ac.uk/entry/S3CXT0 | https://alphafold.ebi.ac.uk/entry/S3CXT0 | https://pubs.acs.org/doi/10.1021/acs.jproteome.9b00740 | N/A |
| A0A7K6M6H4 | Beta barrel TM | https://alphafold.ebi.ac.uk/entry/A0A7K6M6H4 | https://alphafold.ebi.ac.uk/entry/A0A7K6M6H4 | https://pubs.acs.org/doi/10.1021/acs.jproteome.9b00740 | N/A |
| A0A332H2K8 | Globular | https://www.rcsb.org/structure/7TZP | https://www.rcsb.org/structure/7TZP | https://www.biorxiv.org/content/10.1101/2022.11.21.517405v1 | N/A |

### Detailed results

Below we visualize three example proteins Q9UBN4, P42055 and L8XZM1 as displayed in the different tools.

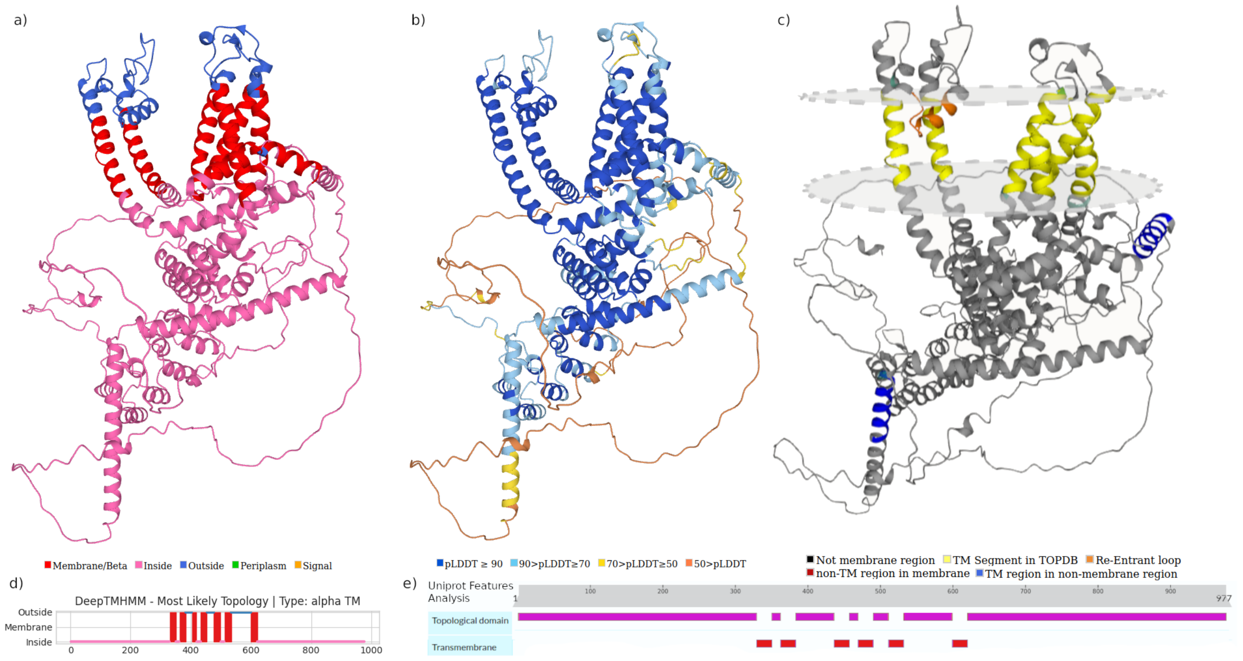


**Figure 1.** Protein Q9UBN4: **a)** structure prediction with DeepTMHMM topology, **b)** structure prediction with AlphaFold Database pLDDT confidence scores, **c)** TmAlphaFold model, **d)** DeepTMHMM most likely topology per residue and **e)** topology retrieved from Uniprot.

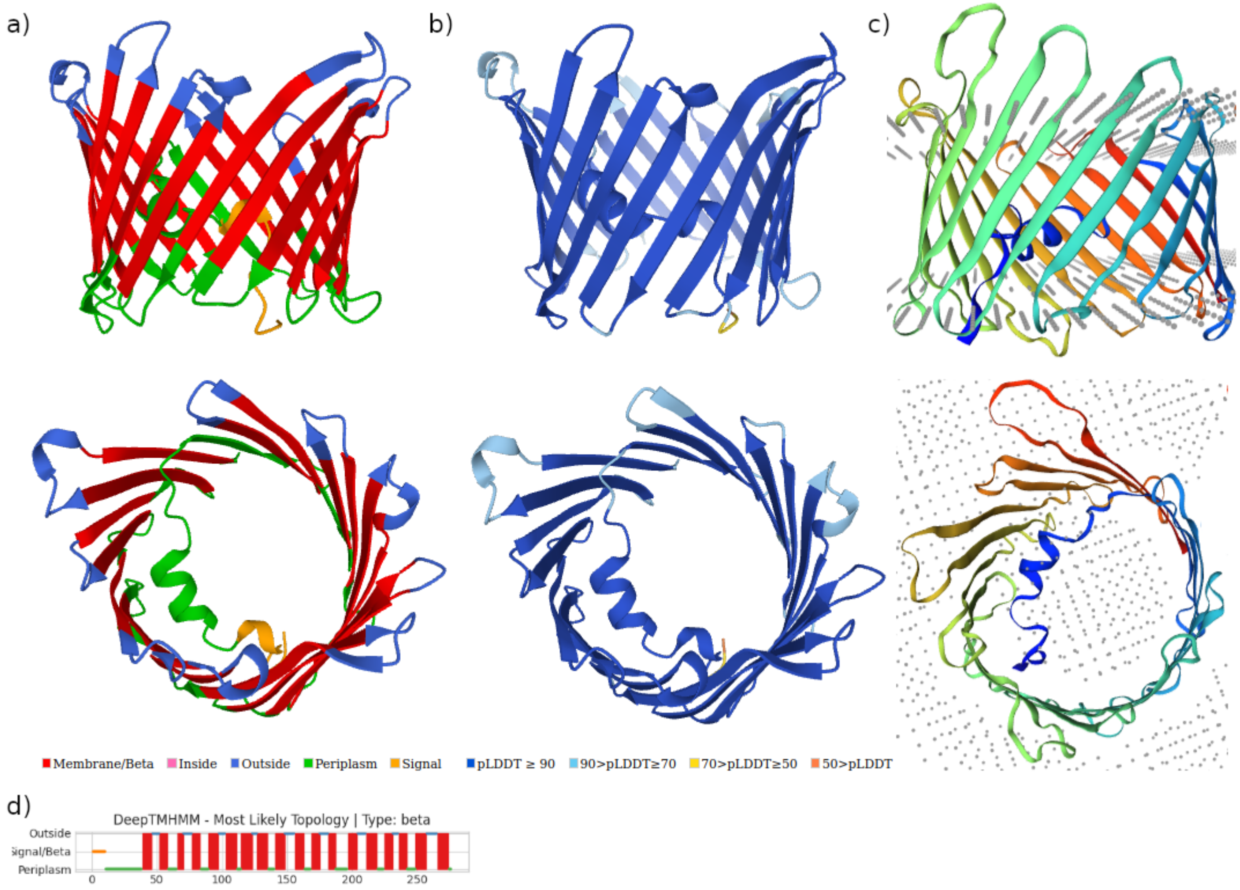


**Figure 2.** Protein P42055: **a)** front (top) and above (bottom) visualizations of the structure prediction with DeepTMHMM topology, **b)** Front (top) and above (bottom) visualizations of the structure prediction with AlphaFold Database pLDDT confidence scores, **c)** Front (top) and above (bottom) visualizations of the protein structure from SWISS-MODEL from https://swissmodel.expasy.org/repository/uniprot/P42055 and **d)** DeepTMHMM most likely topology per residue.

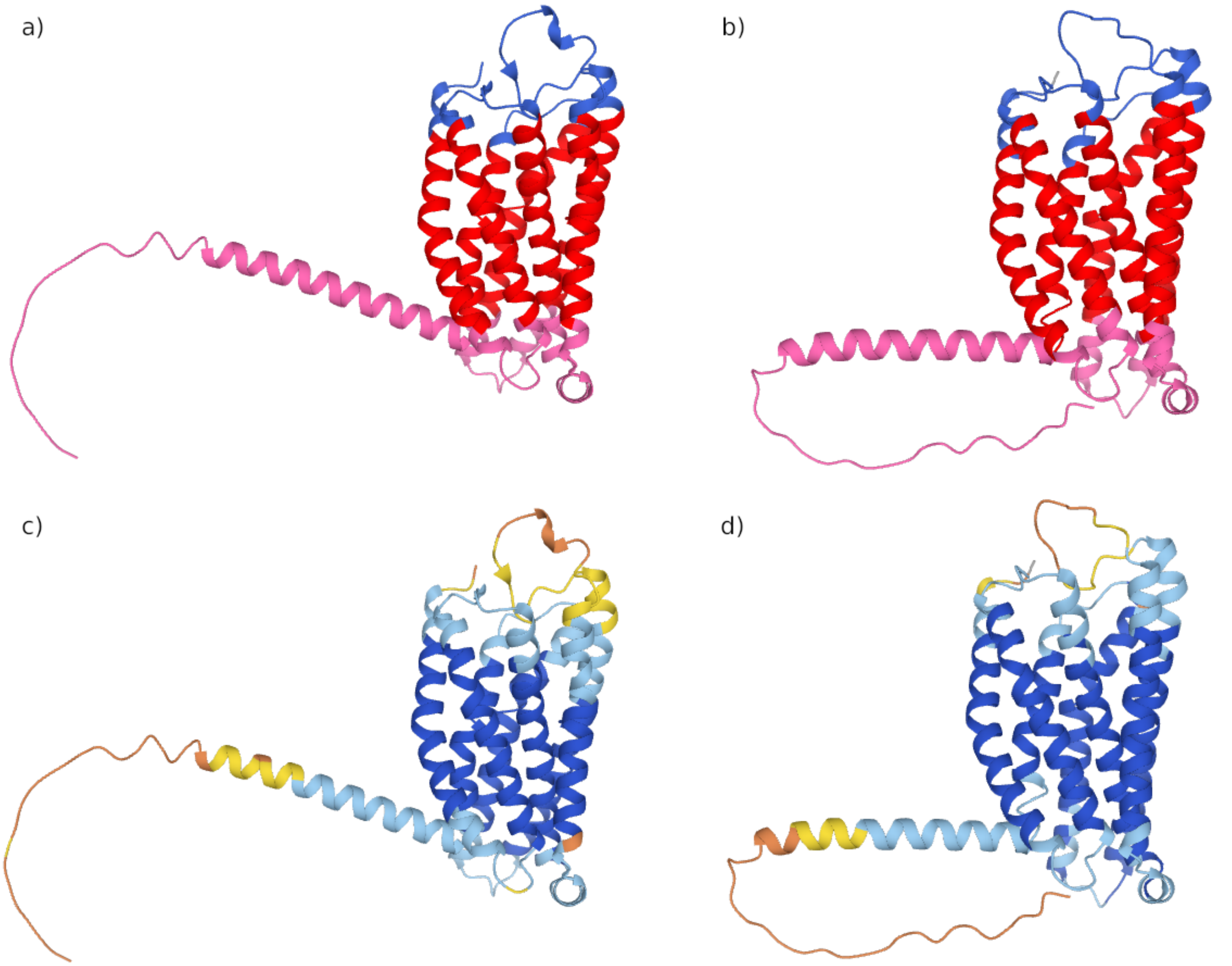


**Figure 3.** Protein L8XZM1: **a)** AlphaFold structure with DeepTMHMM topology, **b)** OmegaFold structure with DeepTMHMM topology, **c)** AlphaFold prediction with pLDDT confidence scores and **d)** OmegaFold prediction with pLDDT confidence scores.
